# Remarked Suppression of Aβ_42_ Protomer-Protomer Dissociation Reaction Elucidated by Molecular Dynamics Simulation

**DOI:** 10.1101/2021.02.12.431048

**Authors:** Ikuo Kurisaki, Shigenori Tanaka

## Abstract

Multimeric protein complexes are molecular apparatuses to regulate biological systems and often determine their fate. Among proteins forming such molecular assemblies, amyloid proteins have drawn attention over a half-century since amyloid fibril formation of these proteins is supposed to be a common pathogenic cause for neurodegenerative diseases. This process is triggered by the accumulation of fibril-like aggregates, while the microscopic mechanisms are mostly elusive due to technical limitation of experimental methodologies in individually observing each of diverse aggregate species in the aqueous solution. We then addressed this problem by employing atomistic molecular dynamics simulations for the paradigmatic amyloid protein, amyloid-β (1-42) (Aβ_42_). Seven different dimeric forms of oligomeric Aβ_42_ fibril-like aggregate in aqueous solution, ranging from tetramer to decamer, were considered. We found additive effects of the size of these fibril-like aggregates on their thermodynamic stability and have clarified kinetic suppression of protomer-protomer dissociation reactions at and beyond the point of pentamer dimer formation. This observation was obtained from the specific combination of the Aβ_42_ protomer structure and the physicochemical condition that we here examined, while it is worthwhile to recall that several amyloid fibrils take dimeric forms of their protomers. We could thus conclude that the stable formation of fibril-like protomer dimer should be involved in a turning point where rapid growth of amyloid fibrils is triggered.

## Introduction

Biological events essential for cell survival are brought about through assembly and disassembly of multiple proteins^1,2^. To understand how these multimeric protein complexes function inside the cell, physicochemical characterization of their formation mechanism is indispensable. It is becoming more feasible to directly examine the microscopic mechanisms due to technical developments in the last two decades^3,4^, whereas their applications to biological systems have been limited so far and the comprehensive knowledge has not yet been obtained satisfactorily.

Among various kinds of assembly of multimeric proteins, amyloid is one of homomeric forms of protein assembly, often referred to as protein aggregates, and has been widely observed in biological systems. Such aggregate products are classified into the two categories, functional amyloid and pathogenic amyloid.^5^ In particular, the latter have drawn much attention because they are supposed as causes of several neurodegenerative diseases, such as Alzheimer and Parkinson diseases.^6^

Since suppression of amyloid formation is considered as practical therapeutic strategy for these serious diseases,^7,8^ molecular mechanisms for their formation have been extensively studied.^9-11^ It is supposed that amyloid fibril formation proceeds via the consecutive three phases. Firstly, monomers assemble to make repertoire of oligomers; a part of oligomers assumes fibril-like, growth-competent aggregates referred to as growth nuclei (lag phase).^12,13^ Secondly, growth nuclei species associate with each other to make larger protofibrils, while they convert natively folded monomers into growth-competent monomers to enhance rapid fibril formation (growth phase). Finally, fibril growth processes are balanced with fibril decomposition processes, then reaching thermal equilibrium and completing fibril formation (plateau phase).

In particular, the progress from lag phase to growth phase plays a critical role in an amyloid fibril formation. Sufficient amounts of growth nuclei species are formed in the lag phase^14^, then triggering amyloid fibril growth^13^. The molecular entity of growth nuclei is regarded as fibril-like aggregates.^13^ Thus clarifying the minimum size of thermodynamically stable fibril-like aggregates, which can be involved in amyloid fibril growth, is a landmark to understand molecular mechanisms for the shift from lag phase to growth phase.

Under these circumstances, an experimental study reported that amyloid protein, amyloid-β (1-42) (Aβ_42_) oligomers can take growth-competent fibril-like forms in the aqueous solution and such protomers bring about the secondary nucleation reactions.^15^ We can suppose that accumulation of such oligomeric Aβ_42_ protomers is one of possible routes leading to conversion from the lag phase to the growth phase. Then understanding microscopic mechanisms of the accumulation process is a promising key to develop new therapeutic strategies for suppressing amyloid formation and thus preventing onsets of several neurodegenerative diseases.

However, it is still challenging to experimentally obtain the microscopic insights into the accumulation processes. This is due to molecular diversity of Aβ_42_ aggregates found in the lag phase^16-19^: it has been technically unfeasible to separately observe and physicochemically characterize each of Aβ_42_ oligomers undergoing aggregation with the atomic resolution.

Then, we address to answer this question with employing atomistic molecular dynamics simulations for oligomeric Aβ_42_ protomers. We here focus on an elementary process of Aβ_42_ protomer accumulation, particularly dimer formation of protomers. Seven Aβ_42_ protomer dimers in aqueous solution are considered as models of Aβ_42_ fibril-like aggregates, where the size of Aβ_42_ protomers ranges from tetramer to decamer. We examined the relationship between thermodynamic stability of Aβ_42_ protomer dimer and the size of protomer. Furthermore, we discussed the association/dissociation mechanism from the structural point of view by testing the hypothesis obtained in our previous study, that Aβ_42_ protomer growth results in suppression of conformational fluctuation such as inter-Aβ_42_ protomer twisting and then thermodynamically stabilizes Aβ_42_ fibril-like aggregates^20^.

We observed kinetic suppression of protomer-protomer dissociation reactions even for Aβ_42_ pentamer dimer. Our observation then suggests that stable formation of oligomeric protomer species is involved in a turning point in Aβ_42_ amyloid fibril formation processes, then giving an important clue toward comprehensive understanding of microscopic mechanisms for shift from the lag phase to the growth phase.

## Materials and Methods

### Setup of amyloid-β (1-42) protomer dimer systems

We used the cryogenic electron microscopy (cryo-EM) structure (PDB entry: 5OQV^21^) to construct amyloid-β (1-42), Aβ_42_, protomer dimer systems; a protomer denotes an Aβ_42_ oligomer composed of Aβ_42_ monomers with growth-competent conformation. Although there is another full-length Aβ_42_ fibril structure (PDB entry: 2NAO^22^), we selected this 5OQV structure by considering the relationship with our earlier study^20^.

Here we consider dimer of *N*-mer Aβ_42_ protomer as the model of fibril-like aggregate, where the value of *N* ranges 4 to 10 (**Figure 1**). Each of the seven models is annotated by Aβ_42_(N:N) or simply N:N, hereafter. Nε protonation state was employed for each of histidine residues, and all carboxyl groups in aspartate and glutamate residues were set to the deprotonated state. Employing each of the seven Aβ_42_(N:N), we prepared seven Aβ_42_ protomer dimer systems, whose annotations and molecular components are summarized in **Table 1**. Since we are interested in relationship between size and thermodynamic stability for these Aβ_42_ protomers, no biological co-solutes were added into aqueous solution except for the counter ions to electronically neutralize these molecular systems. The additional detail for system construction is described in Supporting Information (see **SI-1**).

**Figure 1.**
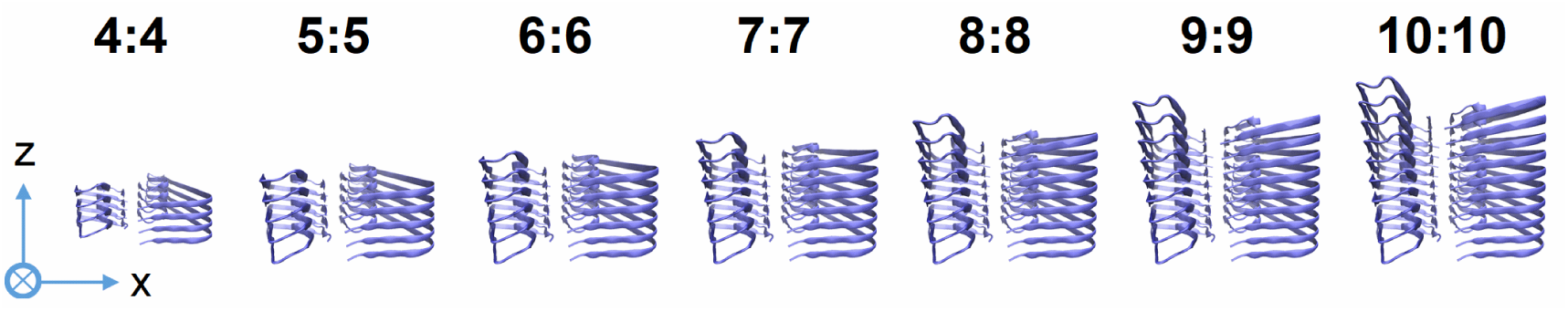
Molecular structures of Aβ_42_ protomer dimers. Each of two integers separated by colon denotes the number of Aβ_42_ monomer in the protomer. The X and Z axes are shown on this plane. The cross in circle denotes Y-axis which directs from this surface to the back.

**Table 1.** Molecular components of Aβ_42_ protomer dimer systems employed for the molecular dynamics simulations.

| size of Aβ<sub>42</sub><br>protomer | number of K<sup>+</sup><br>cations | number of water<br>molecules | box axis length [nm] | | |
| --- | --- | --- | --- | --- | --- |
|  |  |  | x | y | z |
| 4 | 24 | 33406 | 13.8 | 10.4 | 7.8 |
| 5 | 30 | 35679 | 13.9 | 10.5 | 8.3 |
| 6 | 36 | 38153 | 14.0 | 10.6 | 8.8 |
| 7 | 42 | 41156 | 14.1 | 10.7 | 9.3 |
| 8 | 48 | 42498 | 14.1 | 10.7 | 9.7 |
| 9 | 54 | 44855 | 14.2 | 10.8 | 10.2 |
| 10 | 60 | 46828 | 14.2 | 10.9 | 10.6 |

To calculate the forces acting among atoms, AMBER force field 14SB^23^, TIP3P water model^24,25^, and JC ion parameters adjusted for the TIP3P water model^26,27^ were used for amino acid residues, water molecules, and ions, respectively. Molecular modeling of each Aβ_42_(N:N) system was performed using the LEaP modules in AmberTools 17 package^28^.

### Simulation setup

Molecular mechanics (MM) and molecular dynamics (MD) simulations were performed under the periodic boundary condition with GPU-version PMEMD module in AMBER 17 package^28^ based on SPFP algorism^29^ with NVIDIA GeForce GTX1080 Ti. Electrostatic interaction was treated by the Particle Mesh Ewald method, where the real space cutoff was set to 0.9 nm. The vibrational motions associated with hydrogen atoms were frozen by SHAKE algorithm through MD simulations. The translational center-of-mass motion of the whole system was removed by every 500 steps to keep the whole system around the origin, avoiding an overflow of coordinate information from the MD trajectory format. These simulation conditions mentioned above were common in all of the simulations discussed in this manuscript.

### Unbiased molecular dynamics simulation

Following temperature relaxation NVT simulations, 30-ns NPT MD simulations (300 K, 1 bar) were performed and used for following analyses. The system temperature and pressure were regulated with Berendsen thermostat^30^ with a 5-ps of coupling constant and Monte Carlo barostat with attempt of system volume change by every 100 steps, respectively. A set of initial atomic velocities was randomly assigned from the Maxwellian distribution at 0.001 K at the beginning of the NVT simulations. The time step of integration was set to 2 fs. For each Aβ_42_ fibril system, this simulation procedure was repeated thirty times by assigning different initial atomic velocities. The further details are shown in Supporting Information (see **SI-2**).

### Steered and umbrella sampling molecular dynamics simulations

Dissociation processes of Aβ_42_ protomer were described by combining a steered molecular dynamics (SMD) simulation with umbrella sampling molecular dynamics (USMD) simulations. The definitions for reaction coordinates for both SMD and USMD simulations are given in Results and Discussion section.

SMD was employed to dissociate an Aβ_42_ protomer from the remaining part of Aβ_42_(N:N). 0.25-ns SMD simulation was carried out under constant NPT condition (300 K, 1 bar), where the system temperature and pressure were regulated by Langevin thermostat with 1-ps^-1^ collision coefficient, and Monte Carlo barostat with attempt of system volume change by every 100 steps, respectively. The value of reaction coordinate was gradually changed through the SMD simulations by imposing the harmonic potential with the force constant of 4.184 kJ/mol/nm^2^.

Then, certain numbers of snapshot structures were extracted from the SMD trajectory and employed for USMD windows. Following temperature relaxation simulations, several nanosecond NVT USMD simulations (300 K) were performed for each of the USMD windows (**Table S1** in Supporting Information for Aβ_42_ protomer dissociation). The system temperature was regulated using Langevin thermostat with 1-ps^-1^ collision coefficient. Each of the last 1-ns USMD trajectories was used to construct a potential of mean force.

This combined SMD-USMD procedures are repeated eight times for each Aβ_42_ protomer dimer system. Sets of initial atomic coordinates for SMD simulations were randomly selected from the thirty set of unbiased 30-ns NPT MD simulations without allowing duplication. The further details are illustrated in Supporting Information (see **SI-3**).

### Trajectory analyses

Dihedral angle, hydrogen bond (HB) formation and root mean square deviation (RMSd) were calculated with the cpptraj module in AmberTools 17 package^28^. We calculated RMSd to the cryo-EM derived Aβ_42_ protomer dimer structure^21^ using the backbone heavy atoms (i.e., C_*α*_, N, C and O). The geometrical criterion of HB formation is as follows: H-X distance was < 0.35 nm and X-H-Y angle was > 120°, where X, Y and H denote acceptor, donor and hydrogen atoms, respectively.

Each set of USMD trajectories was used to calculate potential of mean force (PMF) with Weighed Histogram Analysis Method (WHAM)^31,32^. Statistical errors of PMF values, *σ*_*PMF*_(*ξ*), were estimated by employing bootstrapped sampling^33^:

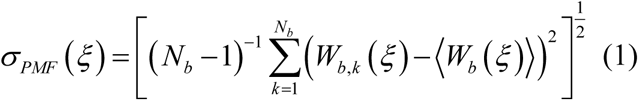

Here, *N*_*b*_, *ξ*, and *W*_*b,k*_ (*ξ*)denote the number of bootstrapped sampling, the reaction coordinate and the value of k^th^ bootstrapped potential of mean force at each point of *ξ*, respectively. ⟨*W*_*b*_ (*ξ*)⟩ is average over all *W*_*b,k*_ (*ξ*), where the value of *N*_*b*_ is set to 200.

Reaction rate, *k*_*TST*_, is estimated by using Eyring’s transition state theory:

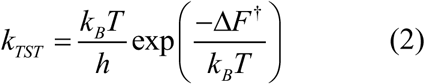

Here, Δ*F* ^†^, *h, k*_*B*_ and *T* denote an activation barrier height, Planck constant, Boltzmann constant and a temperature of system, respectively. Reaction time scale, *τ*_TST_, is defined as the inverse of *k*_*TST*_. Δ*F*^†^ is defined as *F* (*ξ*_0_′) − *F* (*ξ*_0_), where PMF has local minimum at *ξ*_0_, and gradient of PMF turns from positive to negative values at *ξ*_0_′, which is greater than *ξ*_0_. The estimation with employing Eq. 2 is supposed to be an upper bound of the reaction rate (or a lower bound of the reaction time)^34,35^, although this does not essentially change the conclusion we obtained from this study (see related discussion in Results and Discussion section).

Molecular structures were illustrated using Visual Molecular Dynamics (VMD).^36^ Error bars are calculated from standard error and indicate 95% confidence interval if there is no annotation.

## Results and Discussion

### Inter-Aβ_42_ protomer twisting is suppressed through fibril growth

We examined conformational fluctuation of Aβ_42_ protomer dimers under thermal noise by employing each thirty sets of 30-ns unbiased NPT MD simulations. **Figure 2** shows time-course change of averaged RMSd values for Aβ_42_ protomer dimers. For each of the seven systems, it can be considered that the values reach convergence after 20 ns so that we suppose that conformation of each Aβ_42_ protomer dimer is relaxed under aqueous condition. Larger protomer dimers have smaller converged RMSd values, suggesting suppression of conformational fluctuation through increasing sizes of protomers. Aβ(4:4) system shows relatively large RMSd fluctuation compared with the other six systems. Nonetheless, the fluctuation is in magnitude of c.a. 0.2 nm within the time domain and seems smaller than atomic scales, then possibly being insignificant. Accordingly, we employed partial MD trajectories in the period after 20 ns for following analyses.

**Figure 2.**
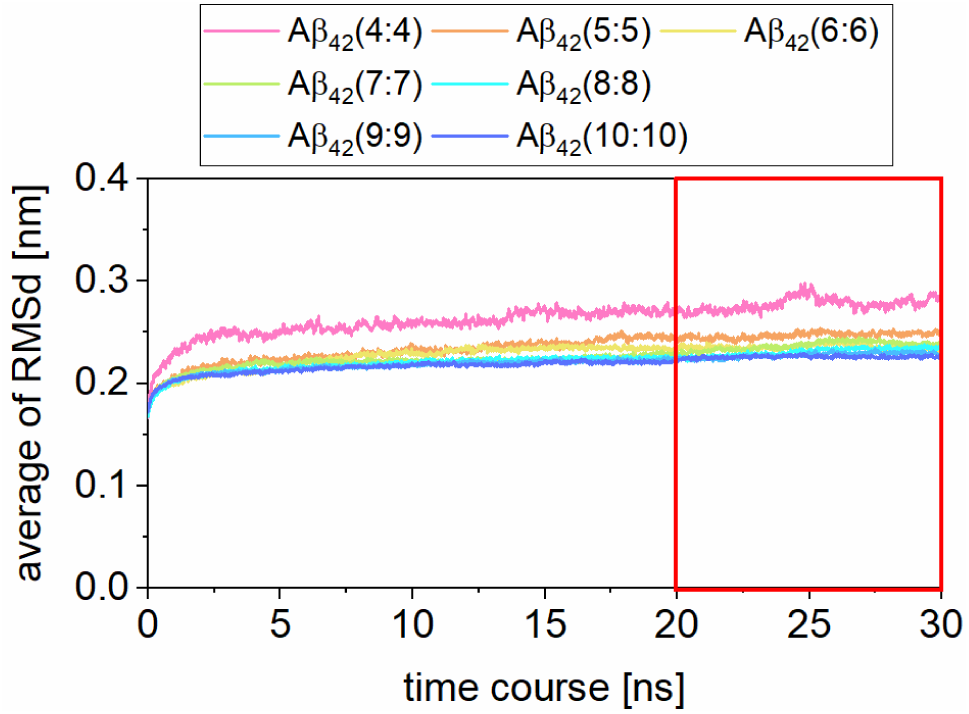
Time-course analyses of root mean square deviation (RMSd). Each result of Aβ_42_ protomer dimer systems is distinguished by color. The time domain supposed as convergence is indicated by the red rectangle.

Recalling the hypothesis about the effect of Aβ_42_ protomer size on the conformational fluctuation of Aβ_42_ protomer dimers, we examined inter-Aβ_42_ protomer twisting angle (*θ*_*T*_) (**Figure 3A**). This *θ*_*T*_ is defined as the dihedral angle in the manner similar to that in our earlier study ^20^. **Table 2** gives statistical analyses for values of *θ*_*T*_ calculated for each of the seven systems. We could not find significant difference in averaged value of *θ*_*T*_ among the seven systems. Meanwhile, standard deviation (S. D.) reflects the effect of protomer size; Aβ_42_(4:4) shows larger S. D. of *θ*_*T*_ than the other systems.

**Figure 3.**
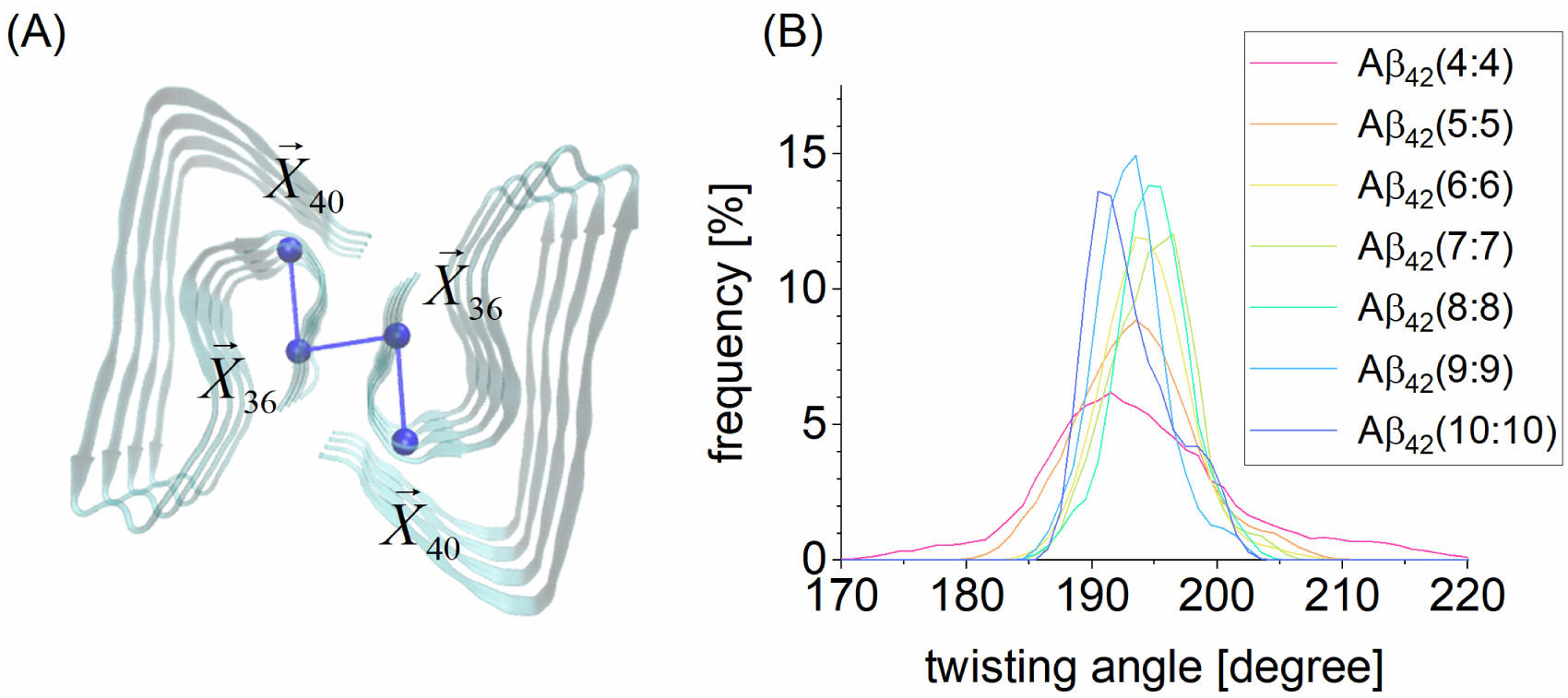
Inter-Aβ_42_ protomer twisting angle. (A) illustration of the twisting angle. (B) twisting angle distributions for each Aβ_42_ protomer dimer system. In panel A, 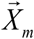 denotes the coordinate averaged for the C_*α*_ atoms of the m^th^ residues in a Aβ_42_ monomer over the protomer (shown with blue ball); Aβ_42_(4:4) is illustrated here as an example. In panel B, each result of Aβ_42_ protomer dimer systems is distinguished by color, and bin width is set to 1°.

**Table 2.**
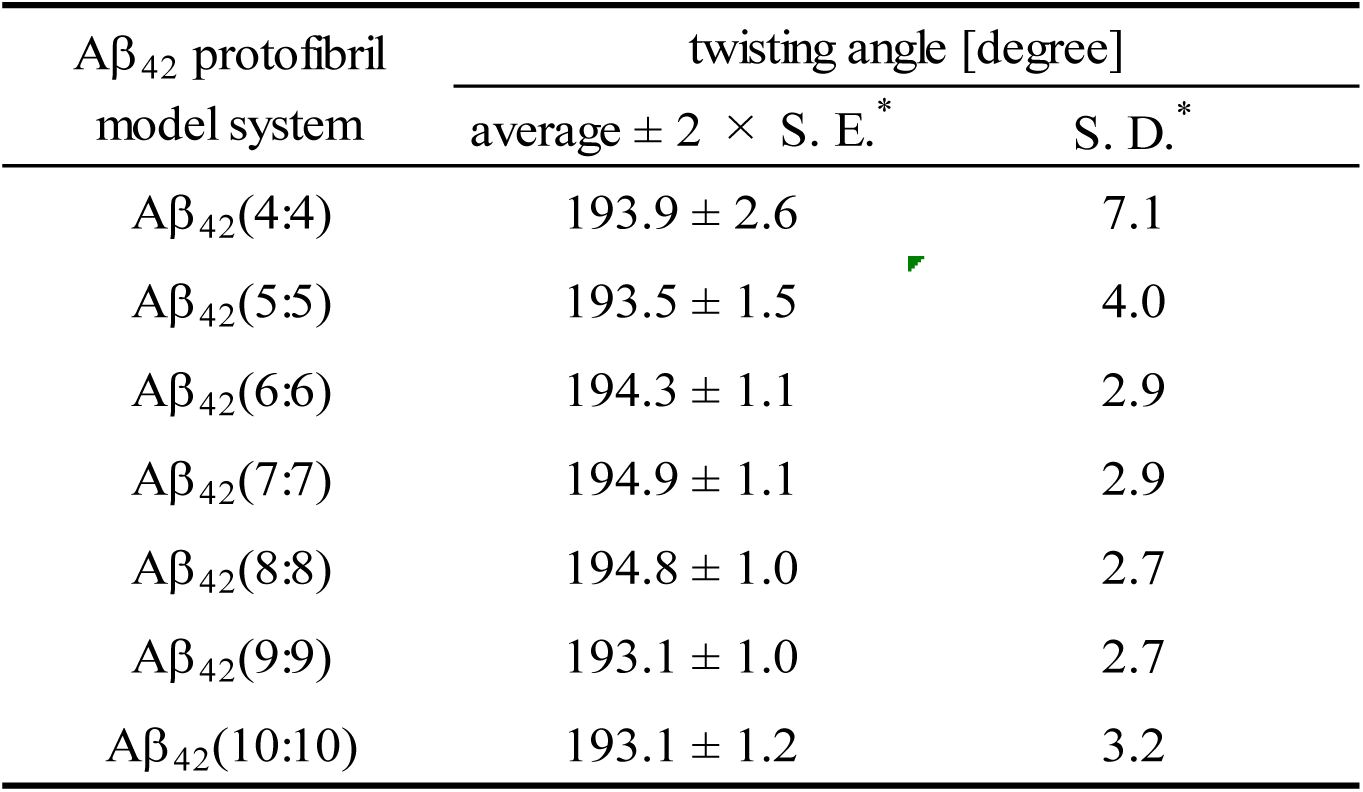
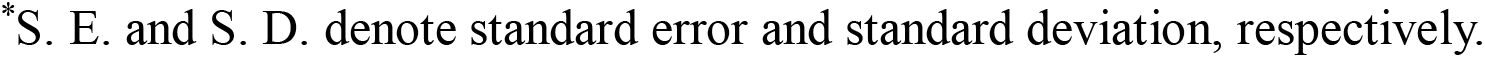
Statistical analyses of inter-Aβ_42_ protomer twisting angle.

| Aβ<sub>42</sub> protofibril<br>model system | twisting angle [degree] |  |
| --- | --- | --- |
| | average $\pm 2 \times$ S. E.* | S. D.* |
| Aβ<sub>42</sub>(4:4) | 193.9 ± 2.6 | 7.1 |
| Aβ<sub>42</sub>(5:5) | 193.5 ± 1.5 | 4.0 |
| Aβ<sub>42</sub>(6:6) | 194.3 ± 1.1 | 2.9 |
| Aβ<sub>42</sub>(7:7) | 194.9 ± 1.1 | 2.9 |
| Aβ<sub>42</sub>(8:8) | 194.8 ± 1.0 | 2.7 |
| Aβ<sub>42</sub>(9:9) | 193.1 ± 1.0 | 2.7 |
| Aβ<sub>42</sub>(10:10) | 193.1 ± 1.2 | 3.2 |

\*S. E. and S. D. denote standard error and standard deviation, respectively.

Dependence of the S. D. on protomer size can be further clarified by showing the shapes of the distributions of *θ*_*T*_ (**Figure 3B**). That for Aβ_42_(4:4) shows longer tails (the S.D. value is 7.0°) than those for other systems. Comparing Aβ_42_(5:5) with Aβ_42_(4:4), the S. D. value shows 1.75-fold decrease, for example. We can find further localization of distributions for larger Aβ_42_ protomer dimers, whose S. D. values are c.a. 3.0°. These observations clearly indicate that increasing the size of Aβ_42_ protomer results in suppression of inter-protomer rotation. This result validates the former part of our hypothesis that Aβ_42_ protomer growth results in suppression of conformational fluctuation such as inter-Aβ_42_ protomer twisting.

### Increase of hydrogen bond formation between the protomers additively changes the height of free barrier of protomer dissociation reaction

We test the latter part of our hypothesis that suppressing conformational fluctuation of Aβ_42_ protomer dimer actually enhances thermodynamic stability of the protomer dimer. A potential of mean force (PMF) was calculated for each of protomer-protomer dissociation reactions, where the distance between centers of mass for each protomer is employed as the reaction coordinates (see **Figure 4A** for the definition).

**Figure 4.**
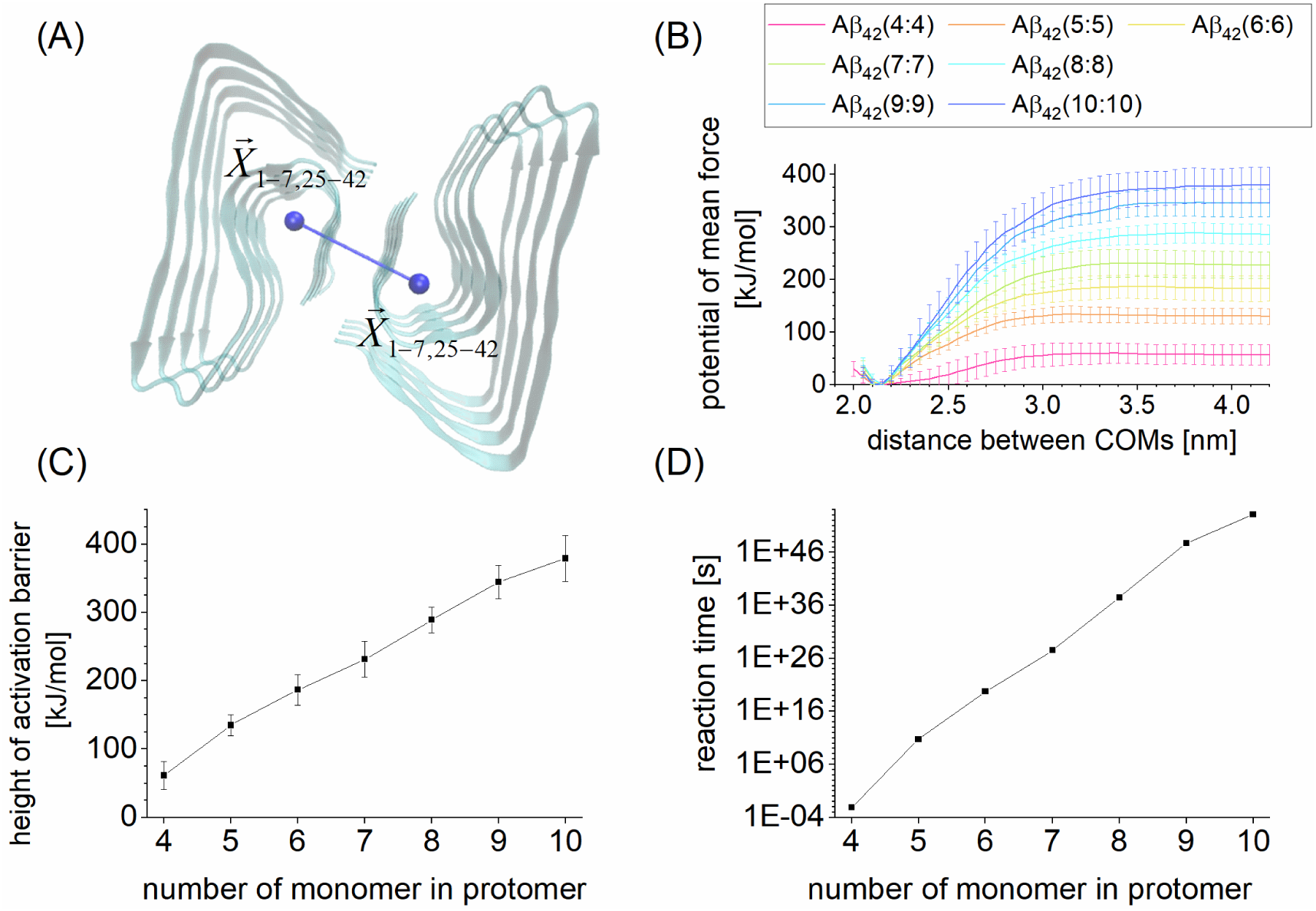
Analyses for Aβ_42_ protomer-protomer dissociation reaction. (A) Illustration of reaction coordinate. (B) Potential of mean forces (PMFs). (C) Height of activation barrier obtained from the PMFs. (D) Reaction time evaluated with Erying’s transition state theory. In panel A,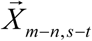 denotes the averaged coordinate (shown with blue ball), which is calculated for the (n-m+1) C_*α*_ atoms of the m^th^, m+1^th^ … n^th^ and s^th^, s+1^th^ … t^th^ residues in the protomer, and the case of Aβ_42_(4:4) is illustrated here as an example. In panel B, each of Aβ_42_ protomer dimer systems is distinguished by color.

As shown in **Figure 4B**, each of the PMFs has one activation barrier toward the dissociation direction, appearing to be uphill. Furthermore, the height of activation barrier clearly increases with the size of protomer, thus corroborating the latter part of our hypothesis (**Figure 4C**).

This change can be explained by considering the number of hydrogen bond (HB) formation between the protomers. **Figure 4C** and **Figure 5A** show the height of activation barrier and the number of inter-Aβ_42_ protomer HB, which is calculated by using both backbone and sidechain atoms in Aβ_42_ monomers, respectively. We can find that both of these two quantities additively change according to protomer size and it thus could be said that increase of the protomer size proportionally contributes to enthalpic stabilization of Aβ_42_ protomer dimers and prevents inter-protomer rotation.

**Figure 5.**
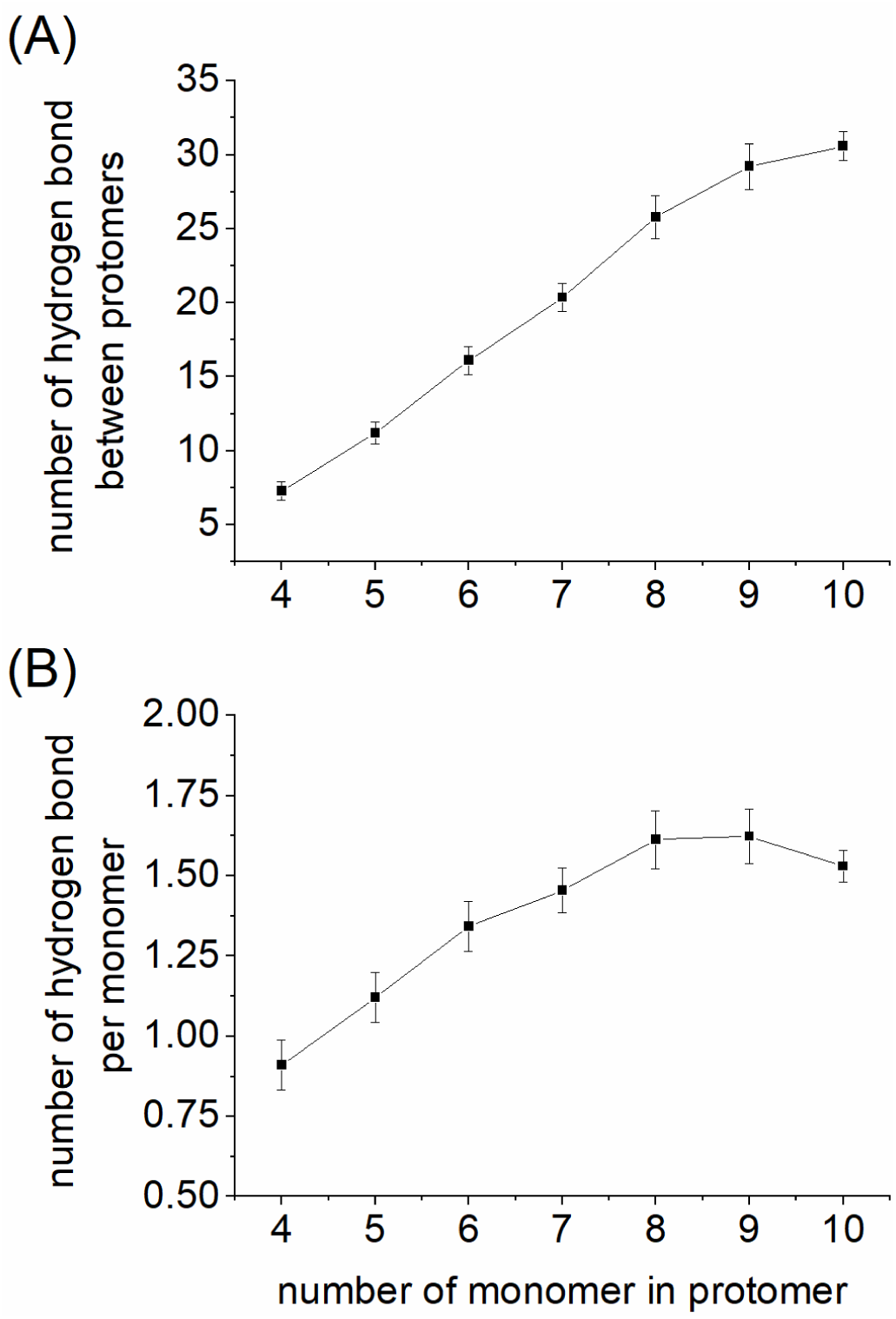
Hydrogen bond (HB) formation between Aβ_42_ protomers. (A) whole number of the HB. (B) the number of HB divided by the number of Aβ_42_ monomer in the Aβ_42_ protomer dimer.

It is worthwhile to note that the number of hydrogen bond formation per Aβ_42_ monomer 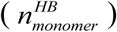 seems to converge up to Aβ_42_(8:8) (**Figure 5B**):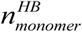 shows apparent monomer size-dependence by Aβ_42_(7:7), while the variable seems to become monomer size-independent from Aβ_42_(8:8). This observation let us suggest that Aβ_42_(8:8) is a boundary where a (relatively) flexible fibril-like aggregate converts into solid Aβ_42_ protofibril structure.

### Protomer-protomer dissociation reaction is remarkably suppressed at the point of Aβ_42_ pentamer formation

Using the heights of activation barriers calculated above, we estimated the time scale of protomer dissociation reactions (*τ*[s]) (**Figure 4D**). From the biological point of view, it is worthwhile discussing the difference between that of Aβ_42_(4:4) and Aβ_42_(5:5), in particular. We can find the critical change of the reaction timescale, due to the exponential dependence on activation barrier height in Eq. 2. According to our estimations with Eyring’s transition state theory, the value of *τ* for Aβ_42_(4:4) is 8.8 ms which falls into timescales of cellular processes, while that for Aβ_42_(5:5), 1680 year, is much longer than mean lifetime of human being. This reaction time estimation for Aβ_42_(5:5) straightforwardly denotes that the protomer-protomer dissociation reaction is kinetically suppressed in decomposition processes of Aβ_42_(5:5) and also in those of greater protomer dimers.

It is noted that Eyring’s transition state theory would often underestimate reaction timescales,^34,35^. Oligomeric species smaller than Aβ_42_(5:5), *e*.*g*., Aβ_42_(4:4), may show suppression of inter-protomer dissociation as well. Nonetheless, the observation for suppression of inter-oligomeric Aβ_42_ protomer dissociation remains unchanged essentially, of course.

The suppression of protomer-protomer dissociation would be common in other Aβ_42_ fibril phenotypes and other amyloid fibrils. It is noted that our observation for the Aβ_42_ pentamer dimer is obtained from the specific combination of the Aβ_42_ protomer structure and the physicochemical condition that we here examined. Meanwhile, several amyloid fibrils take dimeric forms of their protomers^37,38^ so that it can be assumed that protomers composed of a sufficient number of the subunits form the fibril-like dimer and show remarkable thermodynamics stability.

We thus suppose that the remarkable suppression of protomer-protomer dissociation via formation of oligomeric fibril-like aggregates is an important turning point where the lag phase moves to the growth phase, even for other amyloid fibril formation processes proceeding under different physicochemical conditions.

## Conclusion

The accumulation of growth nuclei species triggers conversion from the lag phase into the growth phase, whereas the molecular mechanisms are mostly elusive. In this context, we examined an elementary process of the accumulation, the protomer dimer formations, by considering a paradigmatic amyloid protein, Aβ_42_. With examining the effects of the protomer size on thermodynamic stability of the protomer dimers, we clarified that dimer formation of Aβ_42_ pentamer remarkably suppresses the protomer-protomer dissociation.

Recalling that several amyloid fibrils are found in dimeric form of the protomers^37,38^, we then suppose that suppression of the reaction pathway in the aggregate disassembly process is the common step which promotes accumulation of the growth nuclei species such as oligomeric protomers. Since protein aggregate formations are essentially intricate processes characterized by multiple conformations of specific protein and their complex formations, more detailed elucidations of microscopic and macroscopic mechanisms for their association/dissociation processes remain to be performed computationally and experimentally. Meanwhile, the observation we obtained in this study could be a landmark knowledge to provide further physicochemical insights into formation mechanisms of such complicated molecular assemblies.

## Abbreviations

(Aβ_42_): amyloid protein, amyloid-β (1-42)
(cryo-EM): cryogenic electron microscopy
(MM): molecular mechanics
(MD): molecular dynamics
(SMD): steered molecular dynamics
(USMD): umbrella sampling molecular dynamics
(PMF): potential of mean force
(HB): hydrogen bond

## Supporting Information

The Supporting Information is available. Detailed procedures for unbiased MD, SMD and USMD simulations, Figures and Tables for analyses of these simulations.

## Acknowledgements

This work was supported by a Grant-Aid for Scientific Research on Innovative Areas “Chemistry for Multimolecular Crowding Biosystems” (JSPS KAKENHI Grand No. JP17H06353) and MEXT Quantum Leap Flagship Program (Grant No. JPMXS0120330644).

