## Supporting Information for "Remarked Suppression of Aβ_42_ Protomer-Protomer Dissociation Reaction Elucidated by Molecular Dynamics Simulation"

### SI-1 Construction of amyloid-β (1-42) protomer dimer

We used the cryo-electron microscopy (cryo-EM) structure (PDB entry: 5OQV[1]) to construct $A\beta_{42}$ protomer dimer, $A\beta_{42}$(N:N). $A\beta_{42}$(4:4) derived from the cryo-EM structure was used as the initial structure. Larger $A\beta_{42}$(N:N) were generated by superposition of two $A\beta_{42}$(4:4) on the edges of the protomer dimers and following deletion of overlapped monomer pairs. This procedure was repeated to obtain expected size of $A\beta_{42}$ protomer dimer, which was performed by using root mean square fit with cpptraj module in AmberTools[2].

### SI-2 Unbiased MD simulation for protomer dimer system

For an $A\beta_{42}$ protomer dimer system, the atomic coordinates of water and $K^+$ molecules were energetically relaxed by the following molecular mechanics (MM) and molecular dynamics (MD) simulations. In each of the following MM and MD simulations, the atomic coordinates of non-hydrogen atoms in the $A\beta_{42}$ protomer dimer were restrained by the harmonic potential with force constant of 0.4184 kJ/mol/nm$^2$ (corresponding to 10 kcal/mol/Å$^2$) around the initial atomic coordinates.

First, steric clashes in the system were removed by MM simulation, which consists of 1000 steps of the steepest descent method followed by 49000 steps of the conjugate gradient method. Then the system temperature and density were relaxed through the following five MD simulations: NVT (0.001 to 1 K, 0.1 ps) → NVT (1 K, 0.1 ps) → NVT (1 to 300 K, 20 ps) → NVT (300 K, 20 ps) → NPT (300 K, 300 ps, 1 bar).

The first two NVT MD simulations and the other MD simulations were performed using 0.01 fs and 2 fs for the time step of integration, respectively. The first and second NVT MD simulations were performed using Berendsen thermostat[1] with a 0.001 ps of coupling constant. Meanwhile the following three simulations were performed using Langevin thermostat with 1-$ps^{-1}$ of collision coefficient. In the first NVT MD simulation, the reference temperature was linearly increased along the time-course. In the NPT MD simulation, the system pressure was regulated with Monte Carlo barostat, where the system volume change was attempted by every 100 steps. Each set of initial atomic velocities was randomly assigned from the Maxwellian distribution at 0.001 K.

Using each initial atomic coordinates derived from the above relaxation simulation, an $A\beta_{42}$ protomer dimer conformation also was structurally relaxed in aqueous solution through the following 7-step MD simulations: NVT (0.001 to 1 K, 0.1 ps, 0.4184 kJ/mol/$nm^2$) → NVT (1 to 300 K, 0.1 ps, 0.4184 kJ/mol/$nm^2$) → NVT (300 K, 10 ps, 0.4184 kJ/mol/$nm^2$) → NVT (300 K, 40 ps, 0.2092 kJ/mol/$nm^2$) → NVT (300 K, 40 ps, 0.04184 kJ/mol/$nm^2$) → NVT (300 K, 40 ps) → NPT (300 K, 1 bar, 30 ns). The first two NVT MD simulations and the other MD simulations were performed using 0.01 fs and 2 fs for the time step of integration, respectively. In the first two NVT simulations, the reference temperature was linearly increased along the time-course. In the first 5 steps, non-hydrogen atoms in A$\beta_{42}$ protomer dimer were positionally restrained by the harmonic potential around the initial atomic coordinates. In each NVT simulation, temperature was regulated using Langevin thermostat with 1-$ps^{-1}$ collision coefficient. In the last 30-ns NPT simulation, temperature and pressure were regulated by Berendsen thermostat[3] with a 5-ps coupling constant, and Monte Carlo barostat, where system volume change was attempted by every 100 steps, respectively. The initial atomic velocities were randomly assigned from the Maxwellian distribution at 0.001 K. The snapshot structure obtained from the 30-ns NPT MD simulation procedure was employed for the following SMD simulations.

### SI-3 Calculation of potential of mean force

#### SI-3.1 PMF calculation for protomer dissociation

Steered molecular dynamics (SMD) simulations were performed to prepare the initial atomic coordinates for each window of umbrella sampling (US) MD simulations. As the reaction coordinate for protomer dissociation, we consider the distance between center of mass of C$\alpha$ atoms in one pentamer and that in the other (explained in Fig. 4).

The value of reaction coordinate was gradually changed through the SMD simulations by imposing the harmonic potential with force constant of 4.184 kJ/mol/nm$^2$. The target distance of SMD simulations was set to 6 nm. The SMD simulation was executed for 0.25 ns under NPT condition (300 K; 1 bar). The trajectory was recorded every 5-ps interval. Temperature and pressure were regulated using Langevin thermostat with a 1-ps$^{-1}$ of collision coefficient and Berendsen barostat[3] with a 5-ps coupling constant, respectively. A time step of 2 fs was used to integrate Newton's equation of motion. An initial atomic velocities were taken over from the previous MD simulation. Using each SMD trajectory, we prepared 45 snapshot structures for 45 umbrella windows (Table S1), where a value of the reaction coordinate for each snapshot structure ranges from 2 nm to 4.2 nm with interval of 0.05 nm. For each system, this procedure was repeated 8 times and, the derived snapshot structures were employed for the following USMD simulations.

Using each initial atomic coordinates derived from the SMD simulations, we performed the relaxation simulation and the following USMD simulation. The relaxation simulation consists of 5 steps: NVT (0.001 to 1.0 K, 0.1 ps, 4.184 kJ/mol/nm$^2$) → NVT (1.0 to 300 K, 0.1 ps, 4.184 kJ/mol/nm$^2$) → NVT (300 K, 40 ps, 0.4184 kJ/mol/nm$^2$) → NVT (300 K, 40 ps, 0.2092 kJ/mol/nm$^2$) → NVT (300 K, 40 ps, 0.04184 kJ/mol/nm$^2$). The initial atomic velocities were randomly assigned from the Maxwellian distribution at

0.001 K in the first step. Backbone heavy atoms (Cα, C, N, O) of $A\beta_{42}$ protomer dimer were restrained by the harmonic potential around the initial atomic coordinates. In each of the first two NVT-MD simulations, the reference temperature was linearly increased along the time-course. Then, several nano second USMD simulation was executed under NVT condition (300 K). For each system, the USMD simulation time length is determined by evaluating the convergence of PMF (*see* Panels A-G in Fig. S1). The reaction coordinate of these USMD simulations is the same as that for the SMD simulations. Minimum of the harmonic potential and the corresponding force constant for window of each USMD simulation are summarized in Table S1. In an USMD simulation, temperature was regulated using Langevin thermostat with 1-$ps^{-1}$ collision coefficient, and the interatomic distance for reaction coordinate was recorded every 1-ps interval.

Using 8 sets of 45 USMD simulations, we made a complete histogram spanning the reaction coordinate from 2 nm to 4.2 nm. We confirmed 16% or greater overlap between sampling of neighboring USMD windows, indicating accurate construction of potential of mean force by WHAM[4,5]. Then the complete histogram was employed to compute a potential of mean force, where width of bin was set to 0.05 nm.

**Table S1.** Equilibrium positions of biased potentials and corresponding force constants for protomer dissociation umbrella sampling molecular dynamics simulations, common with the seven A$\beta_{42}$ protomer dimer systems.

| equilibrium position of biasing potential [nm] | force constant [kJ/mol/nm$^2$] 0.4184 | equilibrium position of biasing potential [nm] | force constant [kJ/mol/nm$^2$] 0.4184 |
|---|---|---|---|
| 2.00 | ✓ | 3.15 | ✓ |
| 2.05 | ✓ | 3.20 | ✓ |
| 2.10 | ✓ | 3.25 | ✓ |
| 2.15 | ✓ | 3.30 | ✓ |
| 2.20 | ✓ | 3.35 | ✓ |
| 2.25 | ✓ | 3.40 | ✓ |
| 2.30 | ✓ | 3.45 | ✓ |
| 2.35 | ✓ | 3.50 | ✓ |
| 2.40 | ✓ | 3.55 | ✓ |
| 2.45 | ✓ | 3.60 | ✓ |
| 2.50 | ✓ | 3.65 | ✓ |
| 2.55 | ✓ | 3.70 | ✓ |
| 2.60 | ✓ | 3.75 | ✓ |
| 2.65 | ✓ | 3.80 | ✓ |
| 2.70 | ✓ | 3.85 | ✓ |
| 2.75 | ✓ | 3.90 | ✓ |
| 2.80 | ✓ | 3.95 | ✓ |
| 2.85 | ✓ | 4.00 | ✓ |
| 2.90 | ✓ | 4.05 | ✓ |
| 2.95 | ✓ | 4.10 | ✓ |
| 3.00 | ✓ | 4.15 | ✓ |
| 3.05 | ✓ | 4.20 | ✓ |
| 3.10 | ✓ | | |

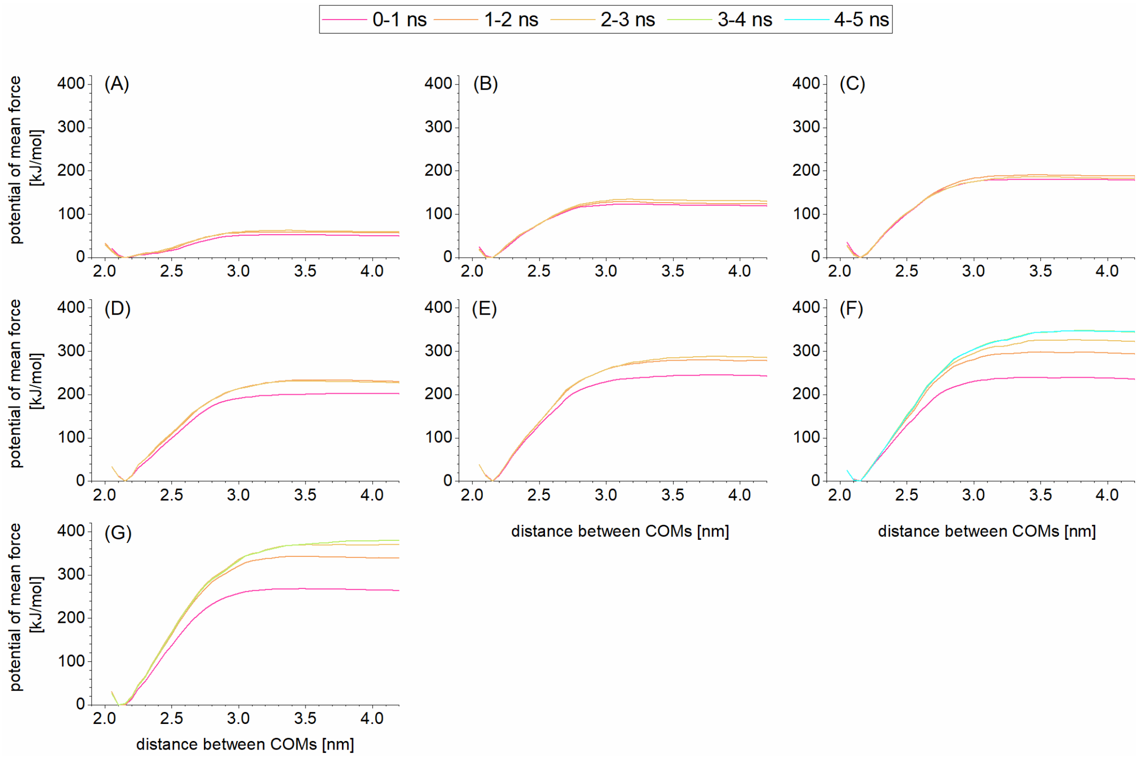


**Figure S1**. Convergence of potential of mean force for protomer dissociation. (A) A$\beta_{42}$(4:4). (B) A$\beta_{42}$(5:5). (C) A$\beta_{42}$(6:6). (D) A$\beta_{42}$(7:7). (E) A$\beta_{42}$(8:8). (F) A$\beta_{42}$(9:9). (G) A$\beta_{42}$(10:10).
